# Hyperrealistic neural decoding: Reconstructing faces from fMRI activations via the GAN latent space

**DOI:** 10.1101/2020.07.01.168849

**Authors:** Thirza Dado, Yağmur Güçlütürk, Luca Ambrogioni, Gabriëlle Ras, Sander E. Bosch, Marcel van Gerven, Umut Güçlü

## Abstract

Neural decoding can be conceptualized as the problem of mapping brain responses back to sensory stimuli via a feature space. We introduce (i) a novel experimental paradigm which uses well-controlled yet highly naturalistic stimuli with a priori known feature representations and (ii) an implementation thereof for HYPerrealistic reconstruction of PERception (HYPER) of faces from brain recordings. To this end, we embrace the use of generative adversarial networks (GANs) at the earliest step of our neural decoding pipeline by acquiring fMRI data as subjects perceive face images synthesized by the generator network of a GAN. We show that the latent vectors used for generation effectively capture the same defining stimulus properties as the fMRI measurements. As such, GAN latent vectors can be used as features underlying the perceived images that can be predicted for (re-)generation, leading to the most accurate reconstructions of perception to date.

## 1 Introduction

Neural decoding can be conceptualized as the inverse problem of mapping brain responses back to sensory stimuli via a feature space [21]. Such a mapping can be modeled as a composite function of linear and nonlinear transformations (Figure 1). A nonlinear transformation models the stimulus-feature mapping whereas the feature-response mapping is modeled by a linear transformation. Invoking this in-between feature space factorizes the direct stimulus-response transformation into two to make it not only data efficient (given that neural data is scarce) but also possible to test alternative hypotheses about the emergence and nature of neural representations of the environment. That is, each stimulus-feature model transforms stimuli to a different set of underlying features to construct candidate feature representations. Each feature-response model then transforms these candidate feature representations to brain responses to test similarity thereof. Feature representations of stimuli are assumed to have a linear relationship with neuroimaging measurements of underlying neural responses such that both capture the same statistical invariances in the environment.

**Fig 1.**
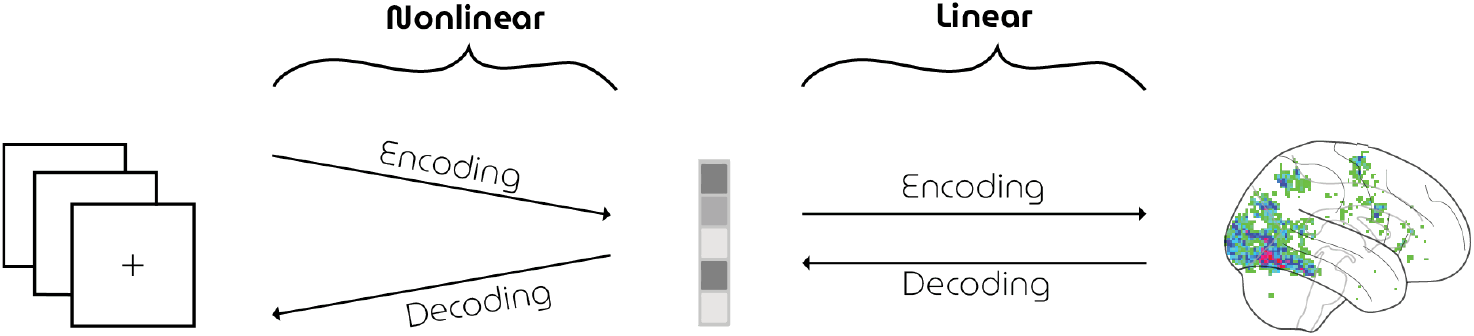
Neural coding. The mapping between sensory stimuli (left) and brain measurements (right) via a feature space (middle). Neural encoding seeks to find a transformation from stimulus to the observed brain response. Conversely, neural decoding seeks to find the information present in the observed brain responses by a mapping from brain activity back to the originally perceived stimulus.

The systematic correspondence between various feature representations of discriminative task-optimized (supervised) convnets and neural representations of sensory cortices are well established [2, 5–7, 14,24]. As such, exploiting this correspondence in neural decoding of visual perception has pushed the state-of-the-art forward [21] such as classification of perceived, imagined and dreamed object categories [9,10], and reconstruction of perceived natural images [16–18] and faces [8,22]. However, unlike their supervised counterparts, more biologically plausible unsupervised deep neural networks have paradoxically been less successful in modeling neural representations [15].

At the same time, generative adversarial networks (GANs) [4] have emerged as perhaps the most powerful generative models to date [1,4,11,12] that can potentially bring neural decoding to the next level. In short, a generator network is pitted against a discriminator network that learns to distinguish synthesized from real data. The goal of the generator is to fool the discriminator by mapping ‘‘latent” vector samples from a given (simple) distribution (e.g., standard Gaussian) to unique data samples such that they appear to have been drawn from the real data distribution. This competition drives the networks to improve in tandem until the synthesized samples are indistinguishable from the real ones. The generator has now learned the unidirectional mapping from latent- to data distribution. This mapping can model the nonlinear feature-stimulus transformation (as defined under neural decoding) where the latent vectors *are* the in-between feature representations underlying the perceived stimuli.

For this reason, GANs have high potential in modeling neural representations but testing this hypothesis is not directly possible because latent vectors cannot be obtained retrospectively; arbitrary stimuli cannot be directly transformed into latent vectors since GANs do not have such an inverse transformation. As a result, the adoption of GANs in neural decoding has been relatively slow since they cannot be readily used for this purpose without resorting to approximate inversion methods (see [17] for such an earlier attempt) unlike those of the aforementioned discriminative convnets. That is, the feature-stimulus transformation entails information loss since the images need to be reconstructed from the predicted feature representations using an approximate inversion network, leading to a severe bottleneck to the maximum possible reconstruction quality (i.e., noise ceiling).

We overcome the aforementioned problem by introducing a very powerful yet simple experimental paradigm for neural (de)coding where participants are presented with synthetic yet highly naturalistic stimuli with known latent vectors. We also present an instance of this paradigm for HYperrealistic reconstruction of PERception (HYPER) which elegantly integrates GANs in neural decoding of faces by combining the following components (Figure 2):

- A pretrained generator network of a progressive growing of GAN (PGGAN) [11] that generates photorealistic face images from latent vectors.
- A new dataset of synthesized face images and whole-brain fMRI activations of two participants.
- A decoding model that predicts latent vectors from fMRI activations which are then fed to the generator for reconstruction.

**Fig 2.**
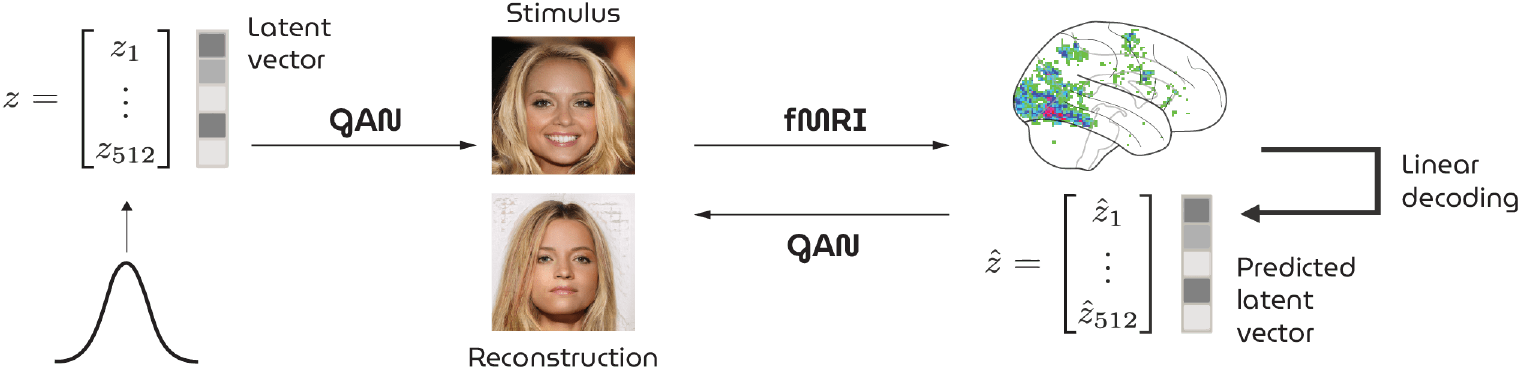
Illustration of the HYPER pipeline. Face images are generated from randomly sampled latent vectors *z* by a GAN and presented as stimuli during brain scanning. A linear model predicts latent vectors *ẑ* for unseen brain responses to feed back to the GAN for reconstruction.

We demonstrate that our approach constitutes a leap forward in our ability to reconstruct percepts from patterns of human brain activity.

## 2 Materials and methods

### 2.1 Neural decoding

The generator of PGGAN is adapted by adding a dense layer at the beginning of the network that performs the response-feature transformation. While the remainder of the network was kept fixed, this response-feature layer is trained by minimizing the Euclidean distance between true and predicted latent vectors (*batchsize* = 30, *lr* = 0.00001, Adam optimization) with weight decay (*alpha* = 0.01) to reduce complexity and multicollinearity of the model.

### 2.2 Datasets

#### 2.2.1 Visual stimuli

High-resolution face images (1024 × 1024 pixels) are synthesized by the generator network of a Progressive GAN (PGGAN) model [11] from latent vectors that are randomly sampled from the standard Gaussian. Each generated face image is cropped and resized to 224 × 224 pixels. In total, 1050 unique faces are presented once for the training set and 36 faces are repeated 14 times for the test set. This ensured that the training set covers a large stimulus space to fit a general face model whereas the voxel responses from the test set contain less noise and higher statistical power.

#### 2.2.2 Brain responses

An fMRI dataset was collected that consists of brain responses to the perceived face stimuli. The fMRI activations (TR = 1.5 s, voxel size = 2 × 2 × 2 mm^3^, whole-brain coverage) of two healthy subjects were measured (S1: 30-year old male; S2: 32-year old male) while they were fixating on a target (0.6 × 0.6 degrees) [20] superimposed on the stimuli (15 × 15 degrees) to minimize involuntary eye movements.

During preprocessing, the brain volumes are realigned to the first functional scan and the mean functional scan, respectively, after which the volumes are normalized to MNI space. A general linear model is fit to deconvolve task-related neural activation with the canonical hemodynamic response function (HRF). Next, we computed the t-statistic for each voxel which was standardized to obtain brain maps in terms of z-scores. In the end, the most-active 4096 voxels on average were selected from the training set to define a voxel mask (Figure 3). Most of these mask voxels are located in the downstream brain regions. Voxel responses from the test set are not used to create this mask to avoid circularity.

**Fig 3.**
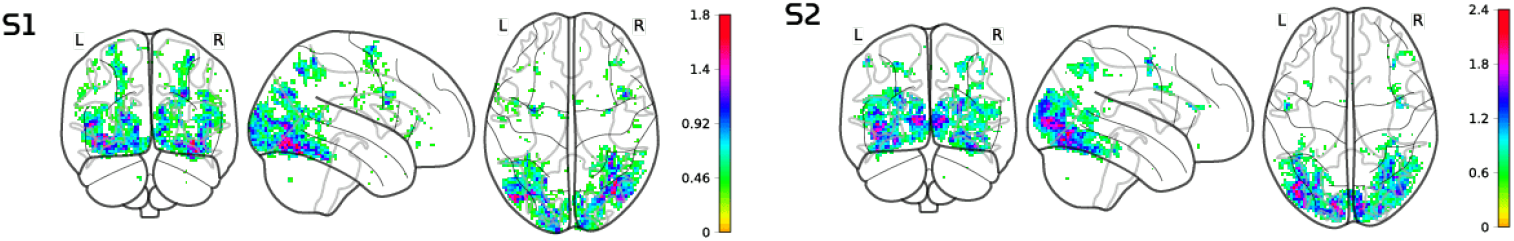
Voxel masks. 4096 most active voxels based on highest z-statistics within the averaged z-map from the training set responses.

The experiment was approved by the local ethics committee (CMO Regio Arnhem-Nijmegen). Subjects provided written informed consent in accordance with the Declaration of Helsinki. The fMRI dataset for both subjects and used models are openly accessible via Github.

### 2.3 Evaluation

Model performance was evaluated in terms of three metrics: latent similarity, feature similarity and structural similarity. First, latent similarity is the Euclidean similarity between predicted and true latent vectors. Second, feature similarity is the Euclidean similarity between feature extraction layer outputs (*n* = 2048) of the ResNet50 model, pretrained for face recognition. Third, structural similarity measured the spatial interdependence between pixels of stimuli and reconstructions [23].

Next, we introduce a new metric “attribute similarity” to assess model performance. Based on the assumption that there exists a hyperplane in latent space for binary semantic attributes (e.g., male vs. female), [19] have identified the decision boundaries for five semantic face attributes in PGGAN’s latent space: gender, age, the presence of eyeglasses, smile, and pose. Attribute scores can be computed by taking the inner product between latent vector and decision boundary. In this way, model performance can be evaluated in terms of these specific visual attributes along a continuous spectrum.

### 2.4 Implementation details

fMRI preprocessing is implemented in SPM12 after which first-order analysis is carried out in Nipy. We used a custom implementation of PGGAN in MxNet together with the pretrained weights from the original paper. Keras’ pretrained implementation of VGGFace (ResNet50 model) is used to evaluate similarities between feature maps of the perceived and reconstructoed images.

## 3 Results

Neural decoding of fMRI measurements via the GAN latent space has resulted in unprecedented stimulus reconstructions. Figure 4 shows arbitrarily chosen but representative examples. The complete set of stimulus-reconstruction pairs can be found in the supplementary materials. Figure 5 illustrates how well the HYPER model captures and decodes attributes by matching polarity and intensity of attribute scores between perceived and reconstructed examples. For most stimulus-reconstruction pairs, the graphs matched in terms of directionality. Correlating observed and predicted feature scores resulted in significant (p ¡ 0.05; Student’s t-test) results for gender, age, eyeglasses and pose, but not for smile (Figure 6).

**Fig 4.**
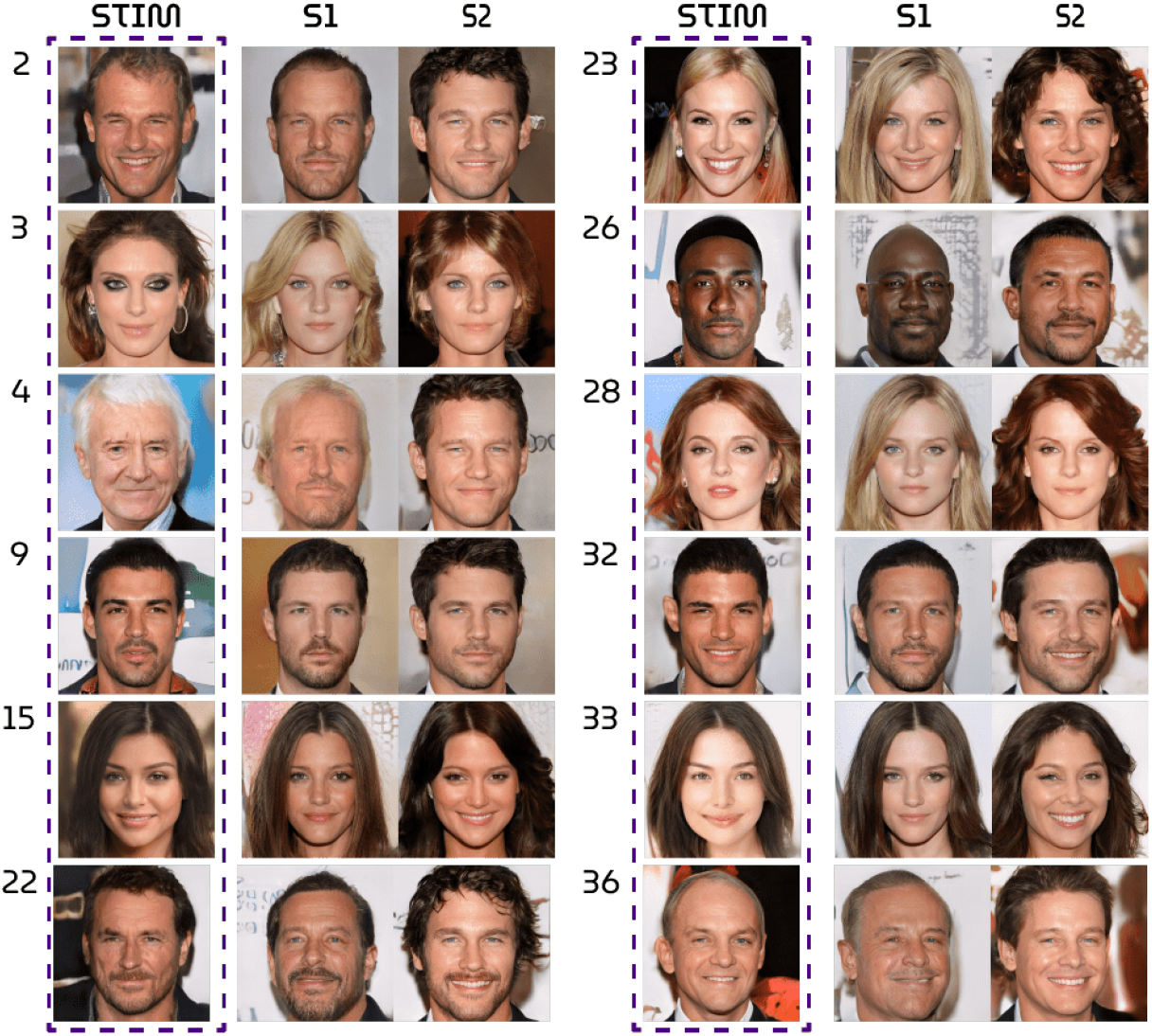
Stimulus-reconstruction examples.

**Fig 5.**
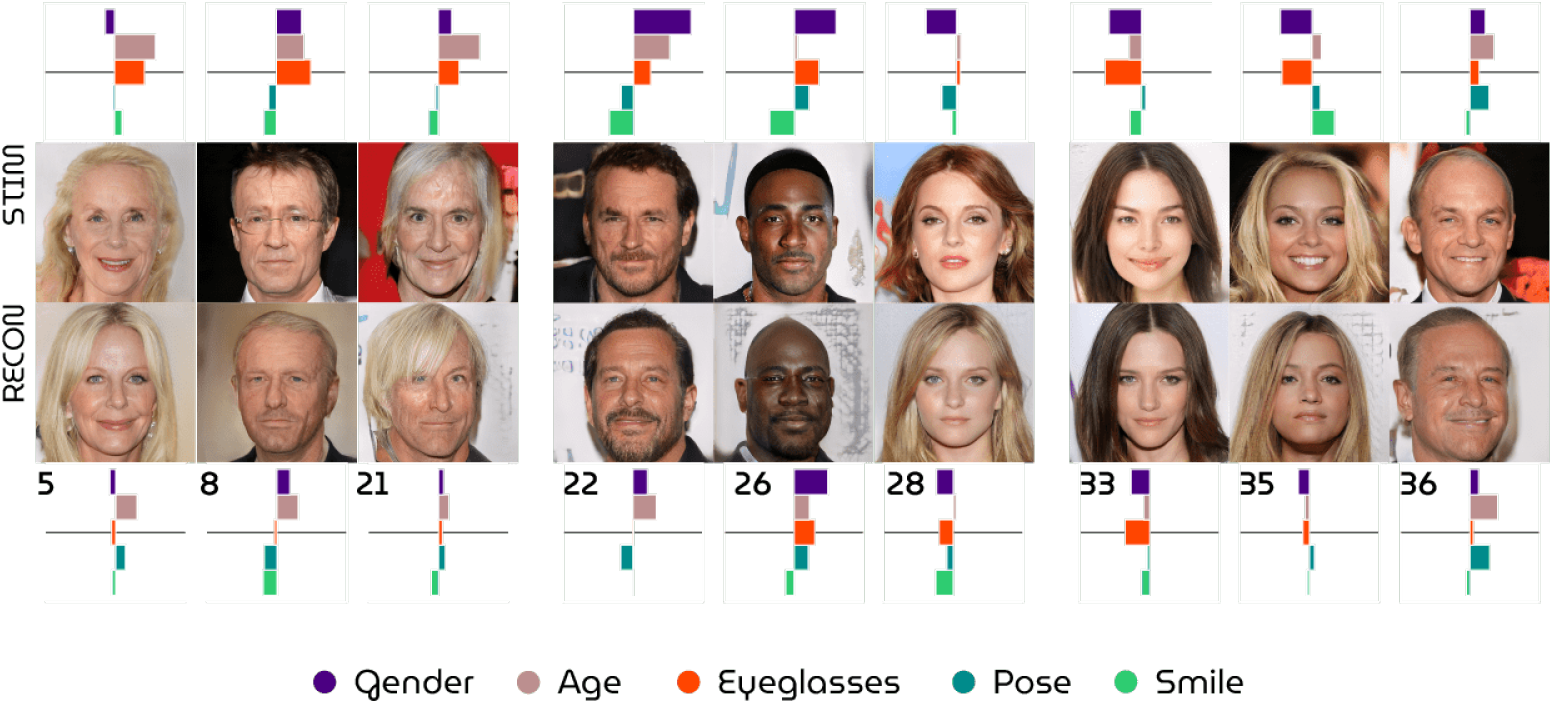
Attribute scores. Stimulus-reconstruction examples from subject 1 together with rotated bar graphs that visualize the attribute scores to demonstrate how attribute similarity can be used to evaluate model performance.

**Fig 6.**
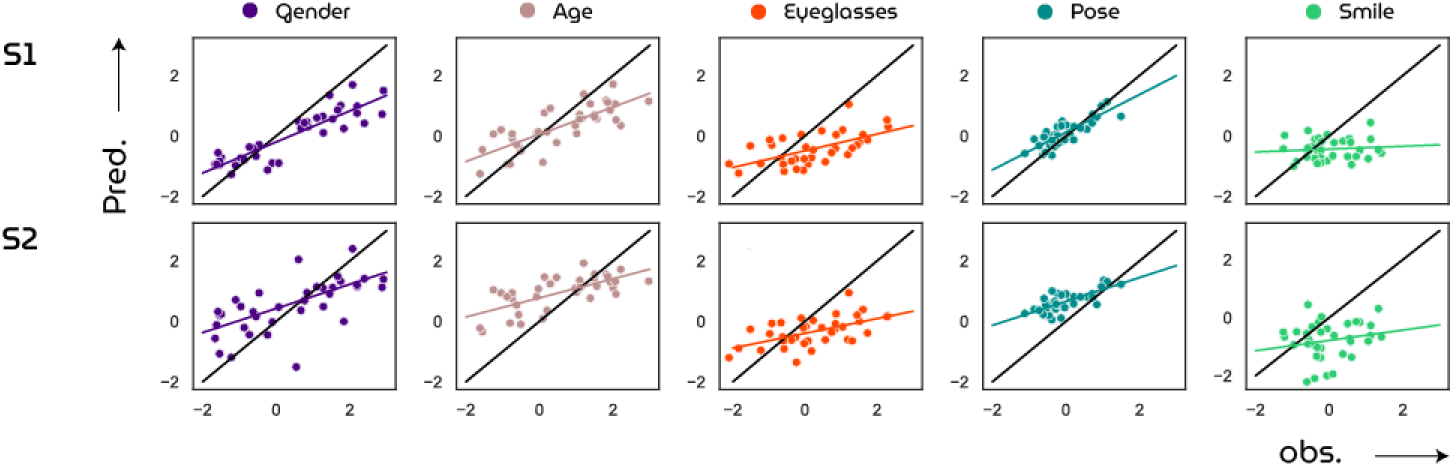
Attribute scores. Significant correlations are found for gender (S1: *r* = 0.89, *p* = 6.55*e* – 13; S2: *r* = 0.20, *p* = 4.20*e* – 10), age (S1: r = 0.78, *p* = 1.70*e* – 08; S2: *r* = 0.69, *p* = 3.91*e* — 06), eyeglasses (S1: *r* = 0.59, *p* = 0.0001; S2: *r* = 0.56, *p* = 0.0003) and pose (S1: r = 0.83, *p* = 4.20*e* — 10; S2: *r* = 0.72, *p* = 4.20*e* — 10), but not for smile (S1: *r* = 0.10, *p* = 0.57; S2: *r* = 0.20, *p* = 0.25).

Next, the performance of the HYPER model is compared to two baseline models that map the brain recordings onto different latent spaces. First, the state-of-the-art VAE-GAN approach [22] predicts 1024-dimensional latent vectors that are fed to the VAE decoder network for reconstruction (128 × 128 pixels). Second, the traditional eigenface approach [3] predicts the first 512 principal components (or “eigenfaces”) and reconstructed face images (64 × 64 pixels) by applying an inverse PCA transform. For a fair comparison, the same voxel masks are used to evaluate all three methods presented in this study without any optimization to a particular decoding approach. All quantitative (Table 1) and qualitative (Figure 7) results showed that the HYPER model outperformed the baselines and had significantly above-chance latent and feature similarity (*p* < 0.001, permutation test) which indicates the probability that a random latent vector or face image would outperform our model predictions.

**Table 1.** Quantitative results. Model performance of the HYPER model compared to the state-of-the-art VAE-GAN approach [22] and the eigenface approach [3] in terms of the feature similarity (column 2) and structural similarity (column 3) between stimuli and reconstructions (mean ± std error). The first column displays latent similarity between true and predicted latents which is only applicable to the HYPER model. For a fair comparison, all images are resized to 224 × 224 pixels, smoothed with a Gaussian filter (kernel size = 3) and backgrounds are removed. Statistical significance of HYPER is evaluated against randomly generated latent vectors and their generated images.

|  |  | Lat. sim. | Feat. sim. | Struct. sim. |
| --- | --- | --- | --- | --- |
| S1 | HYPER | $0.4521 \pm 0.0026$<br>($p < 0.001$; *perm.test*) | $0.1745 \pm 0.0038$<br>($p < 0.001$; *perm.test*) | $0.6663 \pm 0.0115$<br>($p < 0.001$; *perm.test*) |
| | VAE-GAN | - | $0.1416 \pm 0.0025$ | $0.5598 \pm 0.0151$ |
| | Eigenface | - | $0.1319 \pm 0.0016$ | $0.5877 \pm 0.0115$ |
| S2 | HYPER | $0.4447 \pm 0.0020$<br>($p < 0.001$; *perm.test*) | $0.1715 \pm 0.0049$<br>($p < 0.001$; *perm.test*) | $0.6035 \pm 0.0128$<br>($p < 0.001$; *perm.test*) |
| | VAE-GAN | - | $0.1461 \pm 0.0022$ | $0.5832 \pm 0.0141$ |
| | Eigenface | - | $0.1261 \pm 0.0019$ | $0.5616 \pm 0.0097$ |

**Fig 7.**
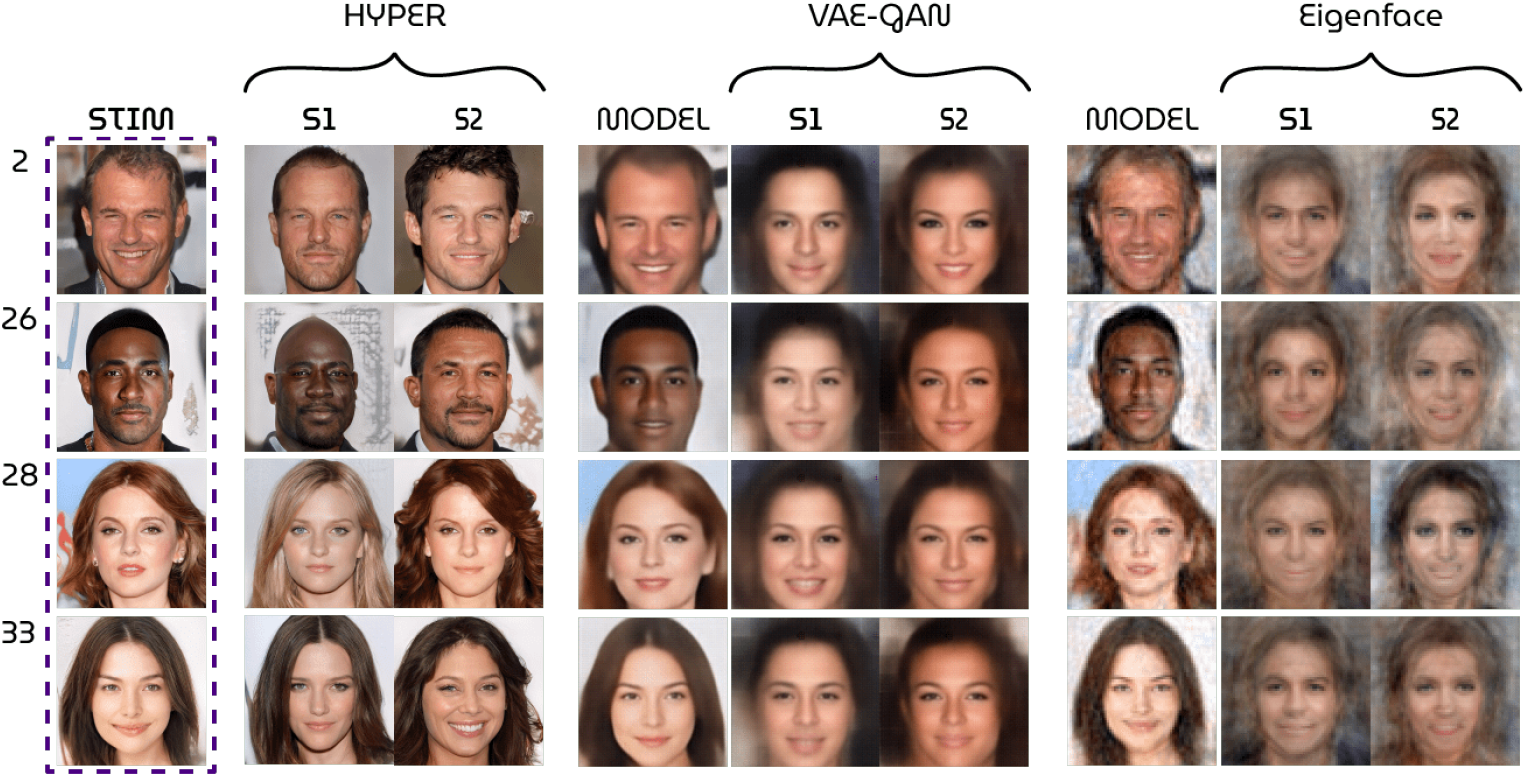
Qualitative results. Model performance of the HYPER model compared to VAE-GAN approach [22] and the eigenface approach [3]. The *model* columns display the best possible results by direct encoding and decoding of the stimulus (i.e., no brain data is used).

## 4 Discussion

The novel experimental paradigm for neural decoding that we introduced uses synthesized stimuli such that the underlying latent/feature representations needed for (re)generation are known a priori. The HYPER model is an implementation of this paradigm which resulted in state-of-the-art reconstructions of perceived faces from brain activations. This indicates that the GAN latent space approximates the neural face manifold and that unsupervised deep neural networks can successfully model neural representations of naturalistic stimuli.

HYPER achieved considerably better reconstructions than the two baselines. It should be noted that the reconstructions by the VAE-GAN approach appear to be of lower quality than those presented in the original study. Likely explanations for this are the differences in dataset size and the voxel selection procedure. Most importantly, we do not attribute the high performance of the HYPER model to the specific type of generative model but instead to the training on synthesized yet photorealistic stimuli (i.e., a GAN is not necessarily better than a VAE-GAN).

### 4.1 Limitations

While HYPER owes its performance to the current advances in generative modeling, it also inherits the limitations thereof in what can and cannot be generated. So far, the generator had to reconstruct faces it already generated before. The next step is verifying whether a decoding model trained on fMRI recordings during perception of synthetic faces generalizes to faces of real people. Latent vectors of real faces are not directly accessible but would also no longer be required when the decoding model has learned to accurately predict them from the synthetic data. It should however be noted that the results of this study are still valid reconstructions of visual perception regardless of the nature of the stimuli themselves.

Reconstructions by HYPER appear to contain biases; the model predicts primarily latent vectors corresponding to young, western-looking faces without eyeglasses as they tend to follow the image statistics of the (celebrity) training set. Also, the PGGAN generator is known to suffer from this problem of *feature entanglement* where manipulating one particular feature in latent space affects other features as well [19]. For example, editing a latent vector to make the generated face wear eyeglasses simultaneously makes the face look older because of such biases in the training data. Feature entanglement obstructs the generator to map unfamiliar latent elements to their respective visual features. It is easy to foresee potential complications for reconstructing images of real faces.

Fortunately, a modified version of PGGAN, called StyleGAN [12,13], is designed to overcome the feature entanglement problem. StyleGAN maps the entangled latent vector to an additional intermediate latent space (thereby reducing feature entanglement) which is then integrated into the generator network using adaptive instance normalization. This results in superior control over the features in the reconstructed images and possibly the generator’s ability to reconstruct unfamiliar features. The generated face photographs by StyleGAN have improved considerably in quality and variation in comparison to PGGAN. Replacing PGGAN with StyleGAN would therefore be a logical next step for studies concerned with the neural decoding of faces.

### 4.2 Future applications

The field of neural decoding has been gaining more and more traction in recent years as advanced computational methods became increasingly available for application on neural data. This is a very welcome development in both neuroscience and neurotechnology since reading neural information will not only help understand and explain human brain function but also find applications in brain computer interfaces and neuroprosthetics to help people with disabilities. For example, extensions of this framework to imagery could make it a preferred means for communication with locked-in patients.

### 4.3 Ethical concerns

Care must be taken as “mind reading” technologies also involve serious ethical concerns regarding mental privacy. Although current neural decoding approaches such as the one presented in this manuscript would not allow for involuntary access to thoughts of a person, future developments may allow for the extraction of information from the brain more easily, as the field is rapidly developing. As with all scientific and technological developments, ethical principles and guidelines as well as data protection regulations should be followed strictly to ensure the safety of potential users of these technologies.

## 5 Conclusion

We have presented a novel experimental framework together with a model for HYperrealistic reconstruction of PERception (HYPER) by neural decoding of brain responses via the GAN latent space, leading to unparalleled stimulus reconstructions. Considering the speed of progress in the field of generative modeling, we believe that this framework will likely result in even more impressive reconstructions of perception and possibly even imagery in the near future.

## A Stimulus-reconstructions

**Fig 8.**
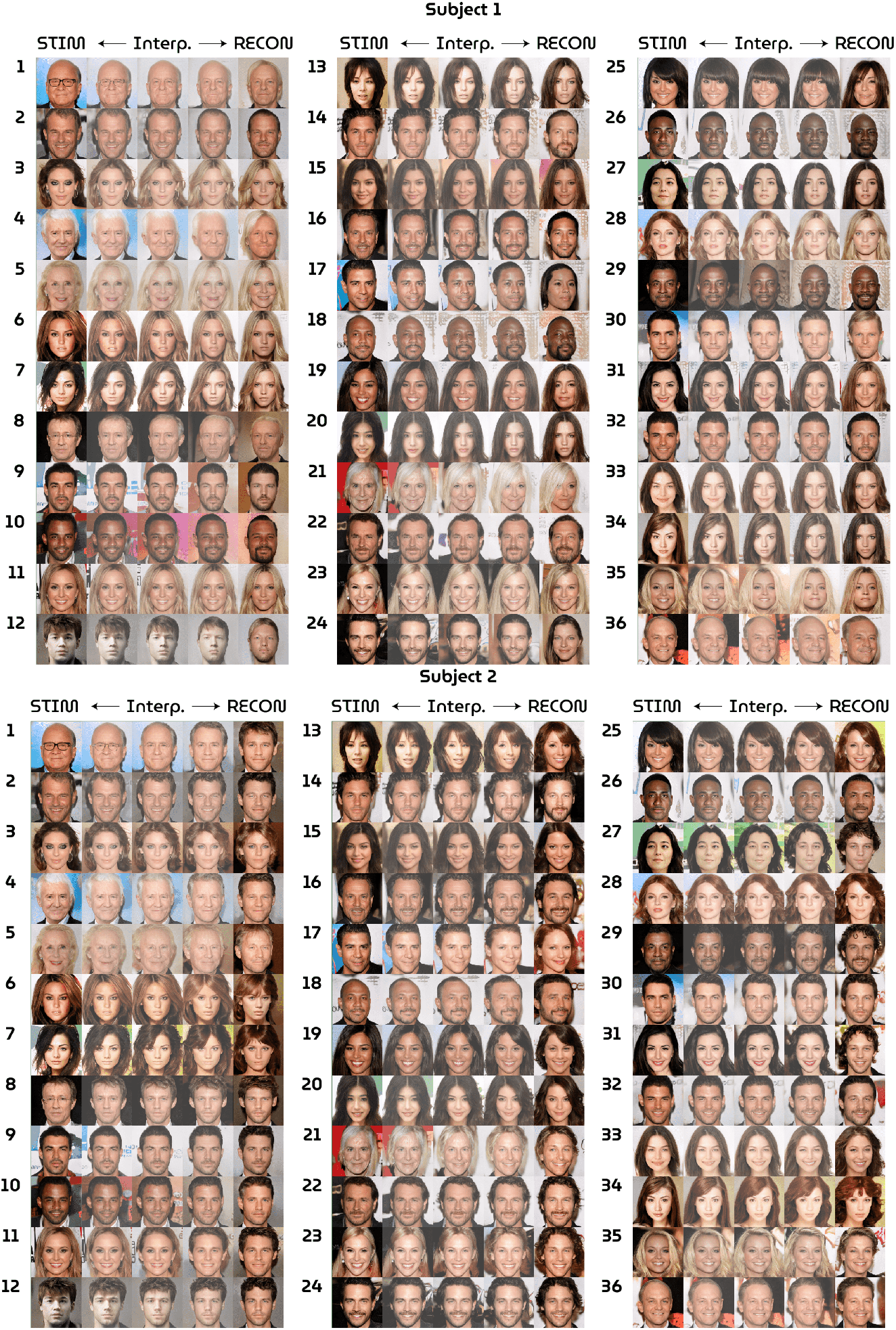
Stimuli (left) and reconstructions (right) for subject 1 and 2. The linear interpolations visualize the distance between predicted and true latent vector that underlie the (re)generated faces. In this way, it is easy to see which features are being retained or change.

## B Github Repository

https://github.com/neuralcodinglab/HYPER

## Notes

### Competing Interest Statement

The authors have declared no competing interest.

## References

[1] Andrew Brock, Jeff Donahue, and Karen Simonyan. Large scale gan training for high fidelity natural image synthesis. arXiv preprint arXiv:1809.11096, 2018.

[2] Charles F Cadieu, Ha Hong, Daniel LK Yamins, Nicolas Pinto, Diego Ardila, Ethan A Solomon, Najib J Majaj, and James J DiCarlo. Deep neural networks rival the representation of primate it cortex for core visual object recognition. PLoS Comput Biol, 10(12):e1003963, 2014.

[3] Alan S Cowen, Marvin M Chun, and Brice A Kuhl. Neural portraits of perception: reconstructing face images from evoked brain activity. Neuroimage, 94:12–22, 2014.

[4] Ian Goodfellow, Jean Pouget-Abadie, Mehdi Mirza, Bing Xu, David Warde-Farley, Sherjil Ozair, Aaron Courville, and Yoshua Bengio. Generative adversarial nets. In Advances in Neural Information Processing Systems, pages 2672–2680, 2014.

[5] Umut Güçlü, Jordy Thielen, Michael Hanke, and Marcel Van Gerven. Brains on beats. In Advances in Neural Information Processing Systems, pages 2101–2109, 2016.

[6] Umut Güçlü and Marcel AJ van Gerven. Deep neural networks reveal a gradient in the complexity of neural representations across the ventral stream. Journal of Neuroscience, 35(27):10005–10014, 2015.

[7] Umut Güçlü and Marcel AJ van Gerven. Increasingly complex representations of natural movies across the dorsal stream are shared between subjects. NeuroImage, 145:329–336, 2017.

[8] Y. Güçlütürk, U. Güçlü, K. Seeliger, S. Bosch, R. van Lier, and M. A. van Gerven. Reconstructing perceived faces from brain activations with deep adversarial neural decoding. Advances in Neural Information Processing Systems, pages 4246–4257, 2017.

[9] Tomoyasu Horikawa and Yukiyasu Kamitani. Generic decoding of seen and imagined objects using hierarchical visual features. Nature Communications, 8(1):1–15, 2017.

[10] Tomoyasu Horikawa and Yukiyasu Kamitani. Hierarchical neural representation of dreamed objects revealed by brain decoding with deep neural network features. Frontiers in computational neuroscience, 11:4, 2017.

[11] Tero Karras, Timo Aila, Samuli Laine, and Jaakko Lehtinen. Progressive growing of gans for improved quality, stability, and variation. arXiv preprint arXiv:1710.10196, 2017.

[12] Tero Karras, Samuli Laine, and Timo Aila. A style-based generator architecture for generative adversarial networks. In Proceedings of the IEEE Conference on Computer Vision and Pattern Recognition, pages 4401–4410, 2019.

[13] Tero Karras, Samuli Laine, Miika Aittala, Janne Hellsten, Jaakko Lehtinen, and Timo Aila. Analyzing and improving the image quality of stylegan. In Proceedings of the IEEE/CVF Conference on Computer Vision and Pattern Recognition, pages 8110–8119, 2020.

[14] Seyed-Mahdi Khaligh-Razavi and Nikolaus Kriegeskorte. Deep supervised, but not unsupervised, models may explain it cortical representation. PLoS Comput Biol, 10(11), 2014.

[15] Seyed-Mahdi Khaligh-Razavi and Nikolaus Kriegeskorte. Deep supervised, but not unsupervised, models may explain it cortical representation. PLoS computational biology, 10(11):e1003915, 2014.

[16] Lynn Le, Luca Ambrogioni, Katja Seeliger, Yağmur Güçlütürk, Marcel van Gerven, and Umut Güçlü. Brain2pix: Fully convolutional naturalistic video reconstruction from brain activity. bioRxiv, 2021.

[17] Katja Seeliger, Umut Güçlü, Luca Ambrogioni, Yagmur Güçlütürk, and Marcel AJ van Gerven. Generative adversarial networks for reconstructing natural images from brain activity. NeuroImage, 181:775–785, 2018.

[18] Guohua Shen, Tomoyasu Horikawa, Kei Majima, and Yukiyasu Kamitani. Deep image reconstruction from human brain activity. PLoS Comput Biol, 15(1):e1006633, 2019.

[19] Yujun Shen, Jinjin Gu, Xiaoou Tang, and Bolei Zhou. Interpreting the latent space of gans for semantic face editing. arXiv preprint arXiv:1907.10786, 2019.

[20] Lore Thaler, Alexander C Schütz, Melvyn A Goodale, and Karl R Gegenfurtner. What is the best fixation target? the effect of target shape on stability of fixational eye movements. Vision Research, 76:31–42, 2013.

[21] Marcel AJ van Gerven, Katja Seeliger, Umut Güçlü, and Yağmur Güclütürk. Current advances in neural decoding. In Explainable AI: Interpreting, Explaining and Visualizing Deep Learning, pages 379–394. Springer, 2019.

[22] Rufin VanRullen and Leila Reddy. Reconstructing faces from fmri patterns using deep generative neural networks. Communications biology, 2(1):193, 2019.

[23] Zhou Wang, Alan C Bovik, Hamid R Sheikh, and Eero P Simoncelli. Image quality assessment: from error visibility to structural similarity. IEEE transactions on image processing, 13(4):600–612, 2004.

[24] Daniel LK Yamins, Ha Hong, Charles F Cadieu, Ethan A Solomon, Darren Seibert, and James J DiCarlo. Performance-optimized hierarchical models predict neural responses in higher visual cortex. Proceedings of the National Academy of Sciences, 111(23):8619–8624, 2014.

